# Integrating Segmental Deuteration iCM-SANS with SAXS and MD for Dynamical Analysis of Multi-domain Proteins

**DOI:** 10.64898/2026.02.25.708105

**Authors:** Aya Okuda, Rintaro Inoue, Minami Kurokawa, Anne Martel, Lionel Porcar, Rikuto Osaki, Kaori Fukuzawa, Kevin L Weiss, Sai Venkatesh Pingali, Reiko Urade, Masaaki Sugiyama

## Abstract

Multi-domain proteins (MDPs) adopt diverse conformations arising from cooperative inter-domain motions, and such dynamics are intimately coupled to their biological functions. Quantitative characterization of these motions is crucial for elucidating their functional mechanisms. Although small-angle X-ray scattering (SAXS) provides information on overall domain arrangement, the limited experimental constraints hinder reliable discrimination of conformational ensembles derived from molecular dynamics (MD) simulations. To address this limitation, complementary experimental constraints that enable to observe domain-selective structural information are required. Inverse contrast-matching small-angle neutron scattering (iCM-SANS), combined with segmental deuteration, enables selective visualization of individual domains and thus provides such complementary information. However, practical strategies for preparing segmentally deuterated MDPs with arbitrary domain labelling have yet to be established. Here, we develop an experimental protocol that integrates controlled protein deuteration with high-efficiency multi-step protein ligation to generate a segmentally deuterated MDP in high yield. The combined use of SAXS and iCM-SANS yields complementary structural constraints that enhance discrimination of MD-derived conformational ensembles. This protocol expands the applicability of segment-selective visualization and also provides an opportunity for high-precision analysis of dynamics in complex MDPs.

**Synopsis:** Segmental deuteration enabled by high-efficiency multi-step protein ligation, combined with inverse contrast-matching SANS and SAXS, provides structural constraints that improve discrimination of molecular dynamics ensembles of multi-domain proteins.

**IMPORTANT:** this document contains embedded data - to preserve data integrity, please ensure where possible that the IUCr Word tools (available from http://journals.iucr.org/services/docxtemplate/) are installed when editing this document.

## 1. Introduction

Multi-domain proteins (MDPs) consist of multiple independently folded structural units (domains) within a single polypeptide chain. When the intervening linkers in MDPs are flexible, domain-domain arrangements can vary widely, leading to diverse conformations. Previous studies have suggested that the MDP functions are not determined solely by the tertiary structures of individual domains, but depend strongly on inter-domain arrangements and their cooperative domain dynamics (Inoue *et al*., 2010; Olsson *et al*., 2010; Roy *et al*., 2016; Vishwanath *et al*., 2018; Inoue *et al*., 2023; Biehl *et al*., 2025). Accordingly, elucidation of MDP functional mechanisms necessitates quantitative characterization of their dynamics under near-physiological solution conditions.

Small-angle X-ray or neutron scattering (SAXS/SANS) techniques using provide spatially and temporally averaged information on molecular shape and structure of the target sample in solution (Guinier *et al*., 1955; Svergun *et al*., 2003; Honecker *et al*., 2022). Taking advantage of this capability, SAXS has been extensively employed as a powerful tool for structural analyses of biomacromolecules in solution and applied to the structural characterization of MDPs (Neylon, 2008; Hou *et al*., 2020; Kawamukai *et al*., 2024). Furthermore, recent advances integrating SAXS with molecular dynamics (MD) simulations have enabled analyses of protein dynamics for less flexible MDPs, such as simple two-domain proteins (Oroguchi *et al*., 2009; de Souza Degenhardt *et al*.; 2021, Shukla *et al*., 2025). In contrast, these approaches still face substantial limitations for more complex MDPs. As increasing the domain number, the internal degrees of freedom arising from domain correlations the conformational degree of freedom expand, resulting in greater conformational diversity and the emergence of more complex structural ensembles. Consequently, conformational ensembles derived from independent MD simulations can produce indistinguishable SAXS profiles despite different domain arrangements (Shevchuk *et al*., 2017). This degeneracy makes it difficult to discriminate appropriate ensembles based solely on SAXS data (Tuukkanen *et al*., 2017), hindering accurate characterization of the dynamics of MDPs. Overcoming this limitation requires integration of SAXS with complementary techniques that directly report on inter-domain arrangements. Such information can subsequently be incorporated as additional experimental constraints.

Neutron exhibits distinct scattering lengths for hydrogen and its isotope deuterium, and this property enables modulation of the difference in scattering length density (SLD) between the solvent and the solute (Heller *et al*., 2010; Krueger *et al*., 2022). Consequently, SANS provides structural information that cannot be obtained from SAXS and has been established as a complementary technique for the structural analysis of biomacromolecules in solution (Svergun *et al*., 1998; Sugiyama *et al*., 2015; Josts *et al*., 2015). Recent advances in protein deuteration techniques, which enable the modulation of SLD of solute, have facilitated the preparation of proteins with controlled degree of deuteration (Okuda *et al*., 2021a). In particular, the SLD of partially deuterated proteins (c.a. 75%) matches that of 100% D^2^O, which has minimal incoherent background scattering. Under this contrast-matching condition, such proteins become “scatteringly invisible” in SANS measurements (Sugiyama *et al*., 2014). Inverse contrast-matching small-angle neutron scattering (iCM-SANS) leverages this principle to achieve selective observation of target components in complexes or multicomponent systems with high signal-to-noise ratios. Indeed, the combined use of SAXS and iCM-SANS data as constraints in structural modeling and/or MD analyses has successfully elucidated both structure and dynamics of protein complex in solution (Yunoki *et al*., 2022). Thus, integration of SAXS and iCM-SANS provides a powerful strategy for high-precision characterization of dynamical analyses of MDPs.

Cooperative dynamics between spatially separated, non-contiguous domains play crucial roles in the functional regulation of MDPs (Yu *et al*., 2010; Liu *et al*., 2022; Agam *et al*., 2024). Investigation of such dynamics requires segmentally deuterated MDPs, where the domains of interest are protonated and the remaining domains are partially deuterated. As a pioneering approach, Sonntag *et al*. demonstrated that a single-step protein ligation enables selective visualization of a contiguous domain using SANS (Sonntag *et al*., 2017); however, extension of this strategy to spatially separated, non-contiguous domains remains unestablished. Preparation of segmentally deuterated MDPs targeting non-contiguous domains requires robust, high-efficiency, multi-step protein ligation strategies beyond current methods. Consequently, combining segmental deuteration of MDPs targeting selective visualization of spatially separated, non-contiguous domains with iCM-SANS enables high-precision dynamical analysis s and provide domain-resolved structural information relevant to functional mechanisms.

In this study, we selected ER-60 (ERp57) (Urade *et al*., 2004; Ruddock *et al*., 2005; Jessop *et al*., 2007; Dong *et al*., 2009, ; PDB ID: 3F8U) as a model MDP. By integrating our developed high-efficiency, multi-step protein ligation strategy (Okuda *et al*., 2023) with a controlled protein deuteration technique (Okuda *et al*., 2021a), we prepared segmentally deuterated samples that facilitate the analysis of inter-domain dynamics between non-contiguous domains. Through iCM-SANS and SAXS measurements, together with correlation analyses based on MD simulations, we propose a methodological protocol that allows high-precision characterization of dynamics of MDPs.

## 2. Materials and Methods

### 2-1. Establishment of expression systems for ER-60 ligation-mutant

In enzymatic protein ligation, an amino acid sequence recognized by the ligation enzyme is typically retained in the final ligation product. In the present system, an NGL sequence remains at the ligation junction. It is therefore essential to verify that this ligation-derived sequence does not perturb the native structure of the target protein. To assess the structural impact of the NGL sequence on full-length ER-60, a corresponding ER-60 variant containing the same substitutions (hereafter referred to as ER-60 ligation-mutant (**a**-**bb’**-**a’**)) was generated in advance. Specifically, four amino acid substitutions (D122N, I124L, R363N, and Y364G) were introduced to create the NGL sequence within the ER-60 backbone.

The DNA plasmid encoding ER-60 ligation-mutant (**a**-**bb’**-**a’**) was generated from the pET-20b(+) plasmid harbouring full-length wild-type human ER-60 (Urade *et al*., 1997) by site-directed mutagenesis PCR using oligonucleotide primer pairs (D122N/I124L: 5′-GCTAATGGACTGGTCAGCCACTTGAAGAAG-3′ and 5′-GACCAGTCCATTAGCAGTCCTAGGTCCATC-3′; R363N/Y364G: 5′-GAAGAACGGCCTGAAGTCTGAACCTATC-3′ and 5′-TTCAGGCCGTTCTTCAGATTGCCATCAAAG-3′) and PrimeSTAR® Max DNA Polymerase (TaKaRa Bio Inc., Japan). The resulting PCR products were directly transformed into *E. coli* TOP10F, and cyclic expression plasmids were obtained. The expression plasmids were transformed into *E. coli* BL21 (DE3) strain (Novagen, Germany).

### 2-2. Preparation of recombinant hydrogenated and deuterated proteins

The recombinant proteins were expressed using *E. coli* expression system. For detailed construct information, see (Okuda *et al*., 2023). The expression of hydrogenated recombinant wild type ER-60, ligation-mutant, **a** and **a’** domain fragments were induced in H_2_O Luria-Bertani (LB) culture containing 0.1 mM isopropyl-β-D-thiogalactoside and corresponding antibiotics (wild-type ER-60 and ligation-mutant: 100 µg/mL Ampicillin, domain fragments: 15 µg/mL kanamycin) at 16°C for 64 h. *E. coli* cells were collected by centrifugation at 6,500 × g for 20 min at 4°C, and disrupted by sonication in 20 mM sodium phosphate-buffered saline (pH 8.0) containing 500 mM NaCl. The homogenate was centrifuged at 26,000 × g for 20 min at 4°C. Soluble proteins in the supernatants were applied to a packed column with Ni Sepharose 6 Fast Flow resin (GE Healthcare, USA). The column was washed with 20 mM sodium phosphate-buffered saline (pH 8.0) containing 500 mM NaCl and 20 mM imidazole. Proteins bound to the resin were eluted with 20 mM sodium phosphate-buffered saline (pH 8.0) containing 500 mM NaCl and 500 mM imidazole. The eluted proteins were subsequently purified by ion-exchange column chromatography on Resource Q (GE Healthcare) with 20 mM Tris-HCl buffer (pH 8.0), followed by gel filtration chromatography on Superdex 200 Increase 10/300GL (GE Healthcare) with 20 mM Tris-HCl buffer (pH 8.0) containing 150 mM NaCl. In the C-side **a’** domains, the His x6-smt3 tag was cleaved by treatment with SUMO protease (LifeSensors, USA) at 4°C overnight. The cleaved tag was removed from the samples by applying them to His60 Ni Superflow Resin (TaKaRa BIO Inc.), followed by size exclusion chromatography (SEC) on Superdex 200 Increase 10/300GL (GE Healthcare) with 20 mM Tris-HCl buffer (pH 8.0) containing 150 mM NaCl. Partially deuterated recombinant domain fragment of ER-60 **bb’** domain was expressed as previously described (Okuda *et al*., 2021a). *E. coli* cells were cultured in 30% D_2_O LB culture solution for 12 h at 37°C. Then, 100 μL of the 30% D^2^O LB culture solution of *E. coli* was added to 5 mL 60% D_2_O LB culture solution and cultured for 12 h at 37°C. The cells in 60% D_2_O LB culture solution were collected by centrifugation and resuspended in 1 L 75% D_2_O M9 medium containing 15 µg/mL kanamycin. Then, the cells were cultured at 37°C until OD_600_ = 0.6. The expression of recombinant ER-60 domain fragment was induced by 0.1 mM isopropyl-β-D-thiogalactoside at 16°C for 64 h. *E. coli* cells were collected by centrifugation at 6,500 × *g* for 20 min at 4°C. The purification procedure was identical to that used for the hydrogenated domain fragments.

### 2-3. Preparation of activation of *oldenlandia affinis* asparaginyl endopeptidase, *Oa*AEP (C247A)

The recombinant *Oa*AEP (C247A) was expressed using *E. coli* expression system. For detailed construct information, see (Okuda *et al*., 2023).The expression of recombinant *Oa*AEP (C247A) were induced in LB broth containing 0.1 mM isopropyl-β-D-thiogalactoside, 15 µg/mL kanamycin, and 12.5 µg/mL tetracycline at 16°C for 64 h. *E. coli* cells were collected by centrifugation at 6,500 × *g* for 20 min at 4°C, sonicated in 20 mM sodium phosphate-buffered saline (pH 8.0) containing 500 mM NaCl, and centrifuged at 26,000 × *g* for 20 min at 4°C. Soluble proteins in the supernatants were applied to a packed column with Ni Sepharose 6 Fast Flow resin (GE Healthcare). The column was washed with 50 mM sodium phosphate-buffered saline (pH 8.0) containing 500 mM NaCl and 20 mM imidazole. The proteins were eluted with 20 mM sodium phosphate-buffered saline (pH 8.0) containing 500 mM NaCl and 500 mM imidazole. To cleave the His x6-smt3 tag, the eluted proteins were treated with SUMO protease (LifeSensors) at 4°C overnight. The cleaved tag was removed from the sample by applying it to His60 Ni Superflow Resin (TaKaRa BIO Inc.). The proteins were subsequently purified by ion-exchange column chromatography on Resource Q (GE Healthcare) with 20 mM Tris-HCl buffer (pH 8.0). To activate its ligation activity of *Oa*AEP (C247A) exposure to a low pH is required (Yang *et al*., 2017). To activate, the purified *Oa*AEP (C247A) (0.5 mg/mL) was dialyzed against 100 mM sodium citrate (pH 4.0) containing 150 mM NaCl and 1 mM EDTA overnight at 4°C. Insoluble proteins generated during dialysis were removed by centrifugation at 6,500 × *g* for 15 min at 4°C. The supernatant was purified by Superdex 75 Increase 10/300GL (GE Healthcare) with 100 mM sodium citrate (pH 4.0) containing 150 mM NaCl.

### 2-4. Ligation reaction of ER-60 domain fragments with *Oa*AEP(C247A)

For the first-step ligation reaction, ER-60 **bb’** and **a’** domain fragments (each at 10 µM) were incubated with 0.2 µM activated *Oa*AEP (C247A) in reaction buffer consisting of 200 mM Tris–HCl (pH 8.0) and 150 mM NaCl for 64 h at 20 °C. The reaction time and temperature were selected based on previously optimized conditions (Okuda *et al*., 2023). After incubation, the reaction mixture was diluted with 2× Laemmli SDS sample buffer (Laemmli, 1970) and heated at 95 °C for 5 min. Samples were analyzed by SDS–PAGE, and gels were stained with Coomassie Brilliant Blue R-250. Band intensities were quantified using ImageJ (National Institutes of Health, USA). Ligation efficiency was calculated as follows:

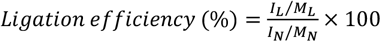

where *I*_L_, *M*_L_, *I*_N_ and *M*_N_ represent the band intensity and molecular weight of the ligation product, band intensity and molecular weight of the N-side fragment in reaction mixture without *Oa*AEP, respectively.

The ligation reaction mixture contained unreacted ER-60 domain fragments, ligation products, and ligation enzyme. Ligation products from the first-step reaction were purified by ion-exchange chromatography on a Resource Q column (GE Healthcare) equilibrated with 20 mM Tris–HCl (pH 8.0). Prior to the second-step ligation, the His_6_–SMT3 tag was removed from the **a** domain fragment using SUMO protease (Ulp1) followed by His60 Ni Superflow resin (TaKaRa Bio Inc.). For the second-step ligation, ER-60 **a** domain and **bb’**–**a’** ligation products (10 µM each) were incubated with 0.2 µM activated *Oa*AEP (C247A) under the same conditions as described above. The resulting ligation products were again purified by Resource Q ion-exchange chromatography (20 mM Tris–HCl, pH 8.0). For final purification of full-length ligated ER-60, the His x6-smt3 tag was removed using Ulp1 and His60 Ni Superflow resin (TaKaRa Bio Inc.), and the product was subsequently subjected to size-exclusion chromatography on a Superdex 200 Increase 10/300 GL column (GE Healthcare) equilibrated with 20 mM Tris–HCl (pH 8.0) containing 150 mM NaCl.

### 2-5. Matrix-assisted laser desorption/ionization-time of flight mass spectrometry (MALDI-TOF MS)

Hydrogenated and deuterated ER-60 domain samples were mixed with a saturated sinapinic acid matrix solution (Bruker Daltonics, Germany) prepared in TA30 (30% acetonitrile in 0.1% TFA aqueous solution) at a sample-to-matrix ratio of 1:9. Aliquots were spotted onto a ground steel MALDI target plate (Bruker Daltonics) and allowed to air-dry and crystallize. Mass spectra were acquired on a microflex LT MALDI–TOF mass spectrometer (Bruker Daltonics) operated in positive-ion mode. External calibration was performed using Protein Standards II (Bruker Daltonics). Spectra were recorded using flexControl and analyzed with FlexAnalysis software (Bruker Daltonics)

### 2-6. SAXS measurement

SAXS measurements were performed using a NANOPIX (Rigaku, Tokyo, Japan). X-rays from a high-brilliance point-focused X-ray generator (MicroMAX-007HF, Rigaku, Tokyo, Japan) were focused with a confocal mirror (OptiSAXS) and collimated with the lower parasitic scattering slit system, “ClearPinhole”. The scattered X-rays were detected with a two-dimensional semiconductor detector (HyPix-6000, Rigaku, Tokyo, Japan) with a spatial resolution of 100 μm. Two sample-to-detector distance (SDD) conditions, 1330 mm and 300 mm, were used to cover a wide *q* range (0.01– 0.70 Å^−1^), where, *q*=(4π/λ)sin(θ/2), with λ (=1.54Å) and *θ* representing the wavelength of X-ray and scattering angle, respectively. The obtained two-dimensional scattering patterns were converted to one-dimensional scattering profiles by radial averaging. Subsequently, one-dimensional scattering profiles were corrected for the intensity of the incident beam and the transmission. Then, the scattering profile of a protein in solution was obtained by subtracting that of the buffer. Finally, the unit of the scatter intensity was converted to an absolute scale by comparison with the scattering intensity of water (*I*_water_ = 1.632 × 10^−2^ cm^−1^). All reductions were processed with SAngler (Shimizu *et al*., 2016).

### 2-7. Analytical ultracentrifugation

The AUC experiments were performed at 60,000 rpm at 25 °C with a sedimentation velocity method using Rayleigh interference optics (ProteomeLab XL-I, Beckman Coulter). The weight concentration distribution *c*(*s*_20,w_) as a function of the sedimentation coefficient was obtained by fitting the time evolution of sedimentation data with Lamm formula (Schuck, 2000) using SEDFIT software (Schuck, 2004). The sedimentation coefficient was also normalized to the value at 20 °C in pure water (*s*_20,w_).

### 2-8. SEC-SANS measurements

SEC-SANS measurements were performed using the D22 instrument at the Institut Laue–Langevin (ILL, Grenoble, France) (Jordan *et al*., 2016; Martel *et al*., 2023) and the Bio-SANS instrument at Oak Ridge National Laboratory (Oak Ridge, TN, USA) (Heller *et al*., 2014; Thomas *et al*., 2024) .

SEC-SANS measurements on D22 were performed using neutrons with a wavelength of *λ* = 6 Å and a wavelength spread of Δ*λ*/*λ* ≈ 10%. By setting the sample-to-detector distance to 8 m for the first detector and 1.4 m for the second detector, the *q* range covered from 0.01 to 0.7 Å^⁻1^. In-line SEC-SANS measurements were performed at the Bio-SANS instrument located at the High Flux Isotope Reactor (HFIR) in the Oak Ridge National Laboratory (ORNL). The main detector array was at 7.0 m from sample, the curved wing detector was fixed at 1.13 m from the sample and rotated to 7.25° from the direct beam, and the curved mid-range detector was fixed at 4m from sample and rotated to 2.7° from the direct beam. Using this 3-detector array configuration, the Q ranges obtained in a single exposure using 6 Å neutrons were 0.007 Å^−1^ < Q < 0.9 Å^−1^. The wavelength spread (Δλ/λ) was 13.2%. The source and sample aperture radii were 20 mm and 7mm, respectively, and separated by 9.7 m. A semitransparent beam trap of borated Aluminum of 38 mm radius was used to acquire transmission and scattering data simultaneously.

In both SEC-SANS measurements, a 500 μL volume of sample at 5 mg/mL was injected onto a Superdex 200 Increase 10/300 GL column (Cytiva). The running buffer consisted of 20 mM Tris–HCl (pH 8.0), 150 mM NaCl, and 1 mM CaCl^2^ in 100% D^2^O. Eluted sample from the SEC column was directly flowed into a flow cell installed on the sample stage of both instruments.

For SEC-iCM-SANS measurements on D22, a UV absorbance detector (280 nm) was aligned at 45° relative to the flow-cell window, enabling real-time monitoring of the elution profile at the cell position. This configuration allowed selective data acquisition at the elution peak of the target fraction. Upon detection of the peak at the cell position, the flow was stopped to retain the eluted peak fraction within the cell for extended acquisition. Data were collected until sufficient counting statistics were achieved. Compared with conventional SEC-SANS measurements under continuous flow, this approach allows longer acquisition times and thereby improves signal-to-noise ratio. Integrated scattering intensity was monitored continuously during measurement to identify the onset of aggregation or degradation. Upon detection of these effects, only data collected before their onset were used to determine the scattering profile of the target fraction.

For Bio-SANS measurements, no UV absorbance detector was available at the flow-cell position. Instead, the UV detector (280 nm) of the HPLC system (ÄKTA pure) located downstream of the SEC column was used to monitor the elution profile. Because sample concentration could not be monitored at the flow cell, the stop-flow strategy used on D22 was not applicable. Therefore, the flow rate was reduced to 0.02 mL min^⁻1^ during peak elution to increase the residence time within the cell, yielding scattering profiles with improved signal-to-noise ratio.

The observed SANS intensity was corrected for background, empty cell and buffer scatterings, and transmission and subsequently converted to the absolute scale using GRASP software (Dewhurst, 2023) or drtsans (Heller *et al*., 2022). Data frames corresponding to the elution peak of the target component were selected, and buffer scattering was subtracted to obtain the scattering profile of the sample.

## 3. Results and Discussion

### 3-1. Preparation of segmentally deuterated ER-60

As illustrated in Figure 1a, ER-60 consists of four thioredoxin-like domains arranged sequentially in an **a**–**b**–**b’**–**a’** architecture. Our previous studies have suggested that changes in the relative orientation between the **a** and **a’** domains could be coupled to its functional regulation (Okuda *et al*., 2021b). In the present study, we prepared a segmentally deuterated ER-60 in which the spatially separated, non-contiguous **a** and **a’** domains were hydrogenated and **b** and **b’** domains were partially deuterated. In this construct, **a** and **a’** domains are “scatteringly visible” and **b** and **b’** domains are “scatteringly invisible” in 100% D^2^O for iCM-SANS measurements (Figure 1b).

**Figure 1.**
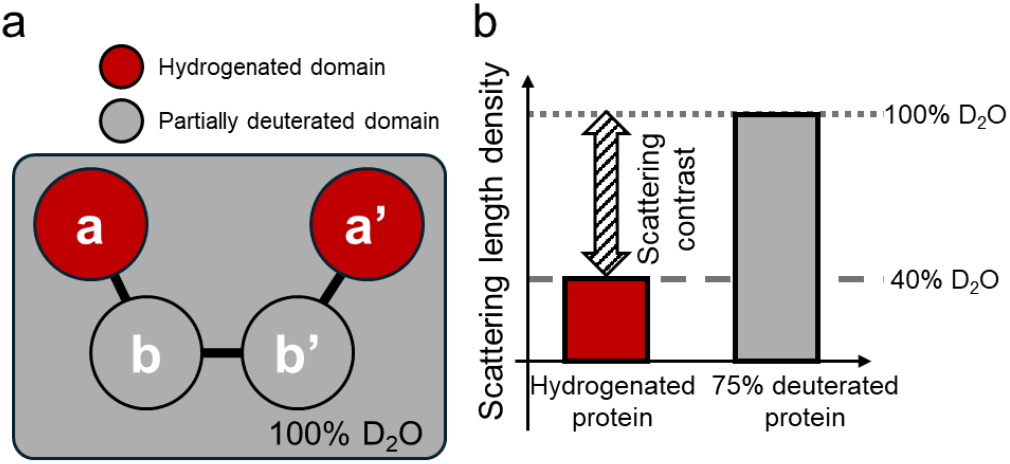
Schematic image of segmentally deuterated ER-60 and scattering length density (SLD) of hydrogenated and partially deuterated proteins. (a) Schematic image of segment deuterated ER-60 comprising partially deuterated **bb’** domains (gray), hydrogenated a (red) and **a’** domains (red). The SLD of partially deuterated **bb’** domains matches that of 100% D_2_O, rendering them “scattering invisible” in SANS. (b) Schematic illustration of scattering length density (SLD) of hydrogenated and partially deuterated proteins. The double-headed arrow indicates the difference of SLD between 100% D_2_O and hydrogenated protein, termed the scattering contrast.

Segmentally deuterated ER-60 was prepared by separately expressing and purifying the domains of interest under different isotopic conditions. The **a** and **a’** domains, designated for selective observation with iCM-SANS, were expressed and purified using an *Escherichia coli* expression system under non-deuterated conditions, whereas the **bb’** domains were expressed and purified under deuterated conditions. These individually prepared domains were subsequently reconstituted into a single polypeptide chain through protein ligation using *Oa*AEP (Figure 2).

**Figure 2.**
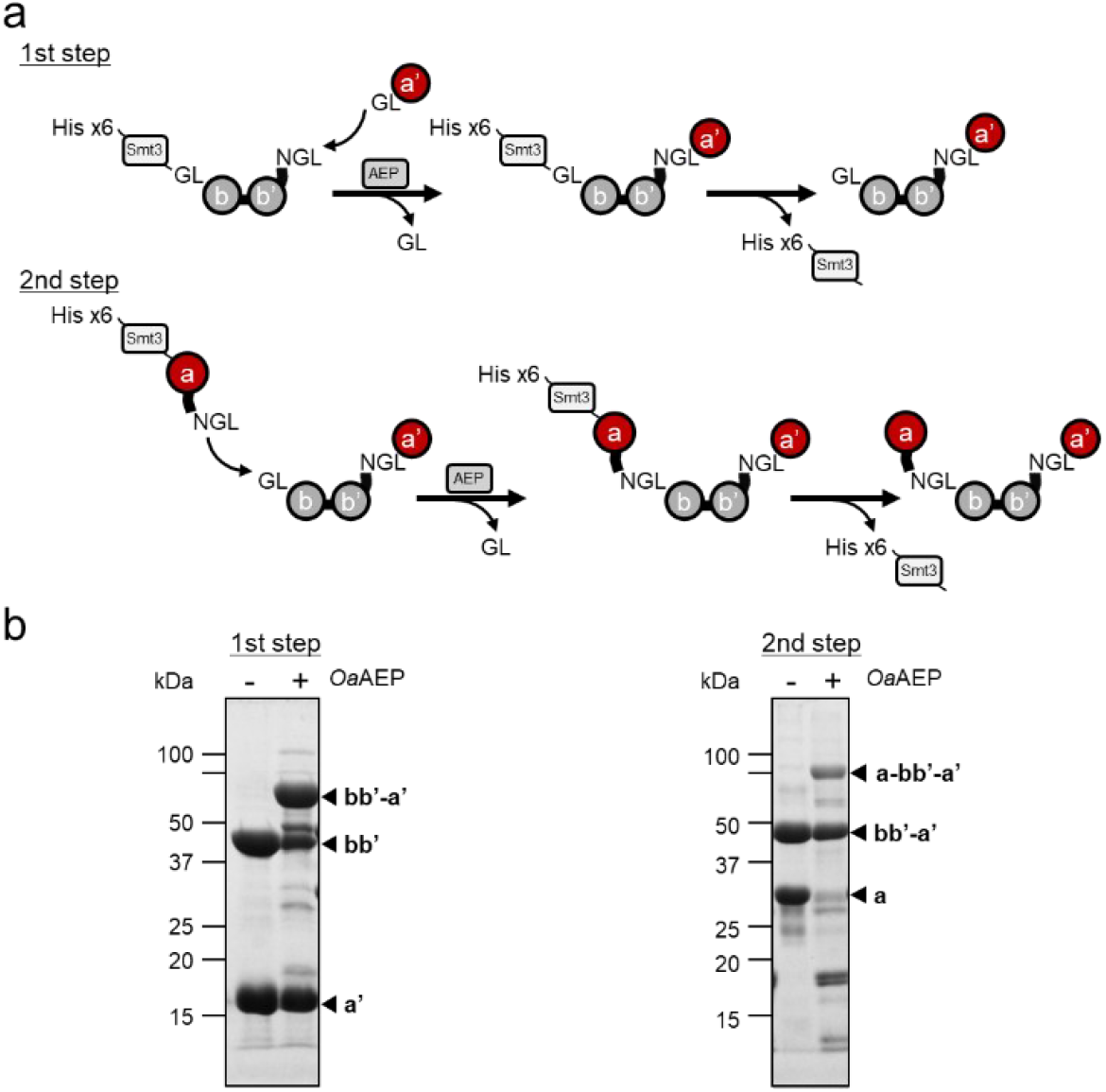
Schematic representation of preparation of segmentally deuterated ER-60 and SDS–PAGE analysis of the first- and second-step ligation reactions.(a) In the first step, the **bb’** and **a’** domains were ligated using *Oa*AEP to generate **bb’**-**a’.** In the second step, the **a** domain was ligated to **bb’**-**a’** to reconstruct full-length ER-60. (b) SDS–PAGE of the two-step ligation reactions. First step: ligation of **bb’** and **a’** domains. Second step: ligation of the **a** domain to **bb’**-**a’**. Lane 1: reaction mixture without *Oa*AEP. Lane 2: reaction mixture with *Oa*AEP.

Using optimized high-efficiency protein ligation with *Oa*AEP, we successfully achieved selective deuteration of specific domains while preserving the native amino acid sequence, except for the minimal *Oa*AEP recognition sequence (NGL) retained after ligation (Figure S1). The degree of deuteration of the **bb’** domain was determined by mass spectrometry (MS) to be 72.5%. This value is in close agreement with the theoretically expected value of 73.9%, calculated from the amino acid sequence to match the SLD of 100% D^2^O (Figure S2). Hereafter, the hydrogenated **a** and **a’** domains are denoted as *h*(**a**) and *h*(**a’**), respectively, while the partially deuterated **bb’** domain is denoted as *pd*(**bb’**). Domains connected by ligation are indicated by hyphens (–). Accordingly, the fully hydrogenated wild-type ER-60 is referred to as *h*(WT), and the hydrogenated ligation-mutant is denoted as *h*(**a**–**bb’**–**a’**).

The ligation efficiencies for the preparation of segmentally deuterated ER-60 are summarized in Table S1. In the first-step ligation between *pd*(**bb’**) and *h*(**a’**), the reaction efficiency reached 62%, which was comparable to that observed for ligation performed exclusively between hydrogenated domains (Okuda *et al*., 2023), indicating that deuteration of the domain had little effect on ligation performance. In contrast, the second-step ligation between *h*(**a**) and *pd*(**bb’**)–*h*(**a’**) exhibited a reduced efficiency of 19%, corresponding to less than half of that observed in the first step. Nevertheless, when 30–50 mg of each domain fragment was prepared, approximately 2.5 mg of the final segmentally deuterated ER-60 product was obtained. By employing optimized *Oa*AEP-mediated ligation conditions, segmentally deuterated MDPs can be prepared for practical SANS experiments. Additionally, the present strategy uniquely enables the preparation of segmentally deuterated ER-60 suitable for selective observation of spatially separated, non-contiguous domains.

### 3-2 Structural characterization of WT, hydrogenated ligation-mutant and segmentally deuterated ER-60 with AUC and SAXS

To evaluate whether the ligation procedure and the associated sequence mutations perturbed the native structure of ER-60, SAXS measurements were performed. In parallel, AUC measurements were conducted using the same samples subjected to SAXS measurements. The AUC analysis revealed the presence of a small fraction of aggregated ER-60 in addition to the monomeric component (Figure S3, Table S2). Because such aggregates could affect the precise analysis of scattering profiles, their contributions must be appropriately removed from observed SAXS profiles. To address this issue, we applied AUC-SAS method (Morishima *et al*., 2020; Morishima *et al*., 2023), which enables extraction of the scattering profile of the target component from the observed SAXS profile of a multi-component system. This approach allowed us to obtain the scattering profile corresponding to monomeric ER-60.

Comparison of the solution structures of wild-type full-length ER-60 (*h*(WT)), the full-length ligation-mutant containing the residual NGL sequence (*h*(**a**–**bb’**–**a’**)), and the segmentally deuterated construct (*h*(**a**)–*pd*(**bb’**)–*h*(**a’**)) revealed no significant differences in their overall domain arrangement in solution (Figure 3, Table 1). These results indicate that neither domain ligation nor segmental deuteration induces substantial perturbations to the overall domain arrangement of ER-60. Furthermore, the present segmental deuteration strategy yielded samples of sufficient quantity and quality for SANS measurements even after multi-step ligation. This ensures that domain-selective structural information obtained from iCM-SANS serves as complementary experimental constraints to the overall structural information derived from SAXS.

**Figure 3.**
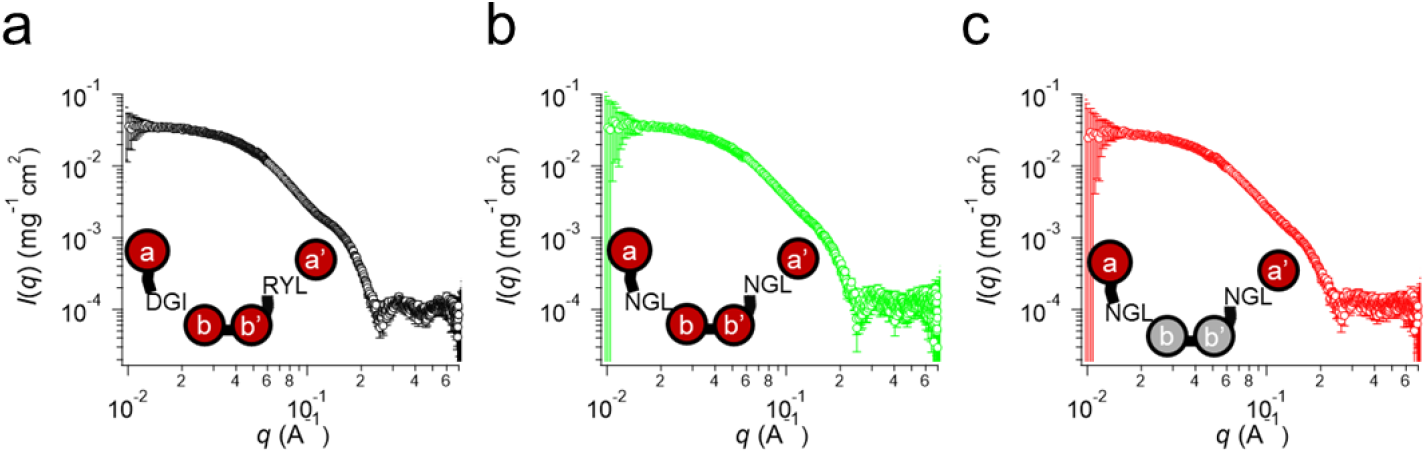
SAXS profiles of ER-60 constructs. SAXS profiles of (a) hydrogenated wild-type ER-60 (*h*(WT)), (b) hydrogenated ligation-mutant (*h*(**a**-**bb’**-**a’**)), and (c) segmentally deuterated ER-60 (*h*(**a**)-*pd*(**bb’**)-*h*(**a’**)). Insets show schematic representations of ER-60 constructs. Red and gray domains indicate hydrogenated and partially deuterated domains, respectively.

**Table 1.**
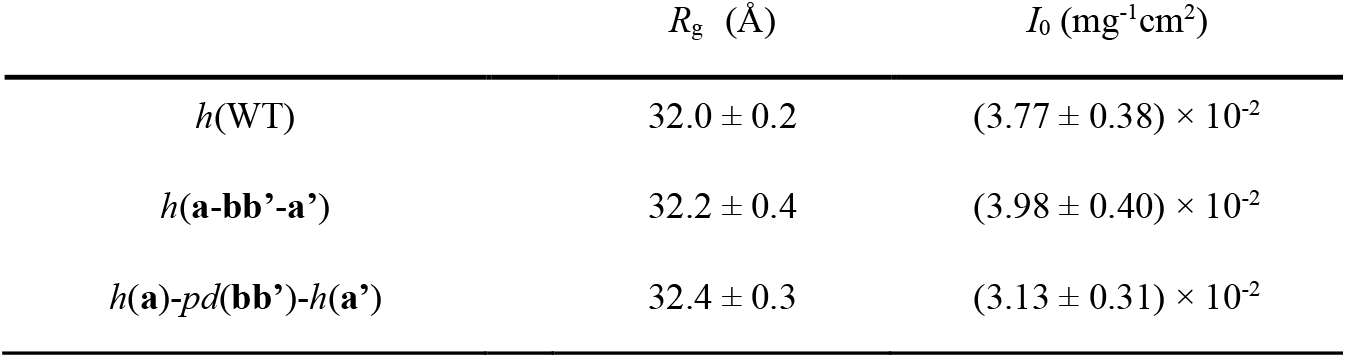
*R*_g_ and *I*_0_ values derived from SAXS profiles of ER-60 constructs.

### 3-3. Inverse contrast-matching SEC-SANS (iCM-SEC-SANS) studies on WT and segment deuterated ER-60

Since aggregates coexisting in solution can affect detailed analysis of SANS scattering profiles as well, we adopted izSEC-SANS, an approach distinct from AUC-SAXS, which also enables extraction of the scattering profile of the target component from a multi-component system.

iCM-SEC-SANS measurements were first performed for the *pd*(**bb’**) in 100% D^2^O (Figure S4a, b). The results demonstrated that the **bb’** domains were effectively contrast-matched, rendering “scatteringly invisible” in iCM-SANS measurements. Accordingly, the scattering contribution from the **bb’** domains can be considered negligible in *h*(**a**)–*pd*(**bb’**)–*h*(**a’**). In other words, only the spatially separated, non-contiguous **a** and **a’** domains are selectively observable from iCM-SEC-SANS.

We then performed iCM-SEC-SANS measurements for *h*(WT) and *h*(**a**)–*pd*(**bb’**)–*h*(**a’**) in 100% D^2^O (Figure 4 and Table 2). SEC chromatograms recorded before SANS (Figure S4c,d), together with scattering-intensity and absorbance traces acquired during iCM-SEC-SANS (Figure S4e,f), confirmed clear separation of aggregated species from the monomeric target component. These results indicate that scattering contributions from aggregates were effectively removed by SEC. For *h*(WT), the overall structure of ER-60 was observed. In contrast, for *h*(**a**)–*pd*(**bb’**)–*h*(**a’**), the scattering profile selectively reflects the relative arrangement of the **a** and **a’** domains. No distinct peak corresponding to the center-of-mass distance correlation between the **a** and **a’** domains was detected. Our previous studies suggested that these domains adopt diverse conformations, and the present results support the notion that they undergo pronounced conformational fluctuations and possess diverse configurations. Such conformational diversity likely accounts for the absence of a clear correlation peak in the scattering profile, indicating that substantial inter-domain dynamics are directly reflected in the scattering profile.

**Figure 4.**
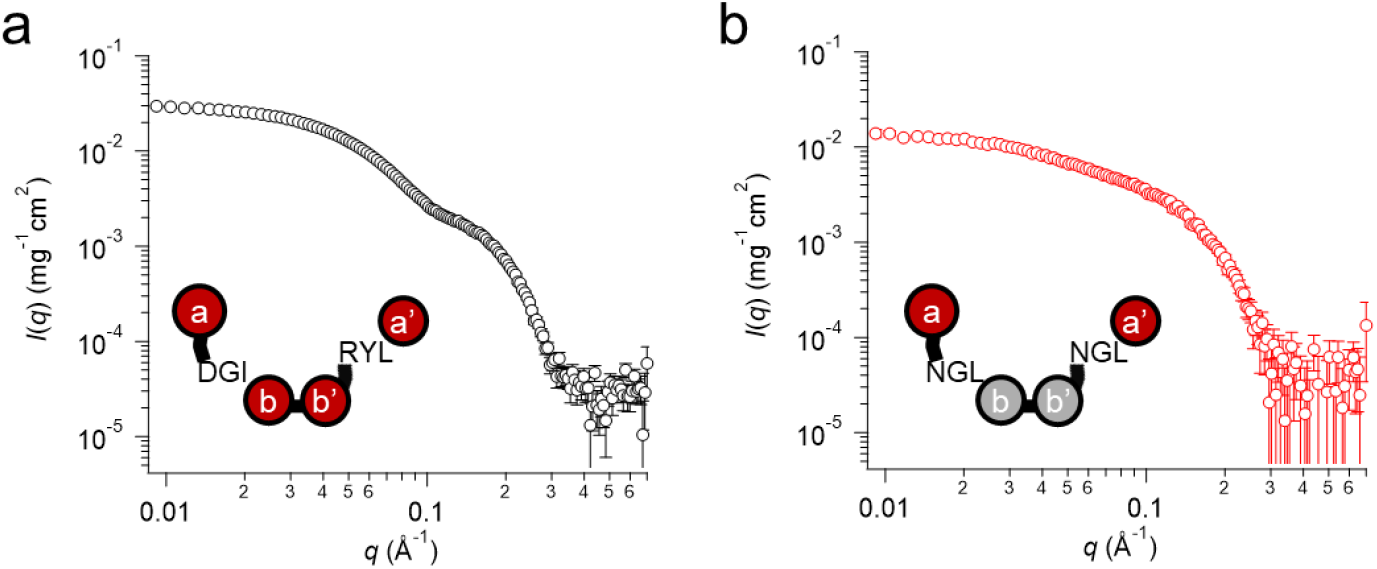
SANS profiles of ER-60 constructs. (a) SANS profile of *h*(WT). (b) SANS profile of *h*(**a**)-*pd*(**bb’**)-*h*(**a’**). Insets show schematic representations of ER-60 constructs. Red and gray domains represent hydrogenated and partially deuterated domains, respectively.

**Table 2.**
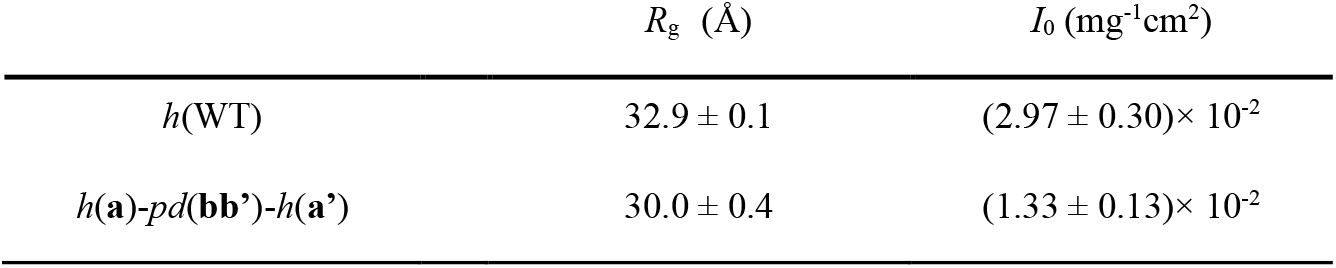
*R*_g_ and *I*_0_ values derived from SANS profiles of ER-60 constructs.

Furthermore, Guinier analysis revealed significant differences in the radius of gyration (*R*_g_) and forward scattering intensity (*I*_0_) between *h*(WT) and *h*(**a**)–*pd*(**bb’**)–*h*(**a’**) (Table 2). The marked decrease in *I*_0_ reflects the reduced total scattering contrast of *h*(**a**)–*pd*(**bb’**)–*h*(**a’**) relative to the 100% D^2^O buffer. Importantly, the observed *I*_0_ ratio between *h*(WT) and *h*(**a**)–*pd*(**bb’**)–*h*(**a’**) agrees with the calculated value. In contrast, the decrease in *R*_g_ might reflect conformations in which the fluctuating **a** and **a’** domains adopt more compact relative arrangements.

These results demonstrate that segmental deuteration combined with iCM-SANS enables experimental visualization of partial structures within MDPs, thereby providing direct structural information on individual domain arrangements and dynamics.

### 3-4. Discrimination of MD trajectories using complementary experimental constraints

The iCM-SANS measurements of segmentally deuterated ER-60 enabled acquisition of domain-specific structural information that cannot be obtained from SAXS alone. However, it remained to be verified whether such partial structural information indeed provides complementary constraints to the SAXS profile and whether it can effectively contribute to the selection of conformational ensembles derived from MD simulations. To examine this point, we performed ten independent 100 ns MD simulations of ER-60. It should be noted that these simulations were conducted solely for proof-of-concept validation of the methodological protocol in this study and therefore may not fully sample the conformational ensemble of ER-60 within this timescale.

From each trajectory, the inter-domain distance between the **a** and **a’** domains and the angle defined by the center-of-mass of the **a, bb’**, and **a’** domains (θ_**a**–**bb’**–**a’**_) were calculated (see Figure 5a). The resulting structural ensembles defined by these two parameters (Figure 5b) revealed that while some trajectories exhibited similar distributions, substantial heterogeneity was observed overall.

**Figure 5.**
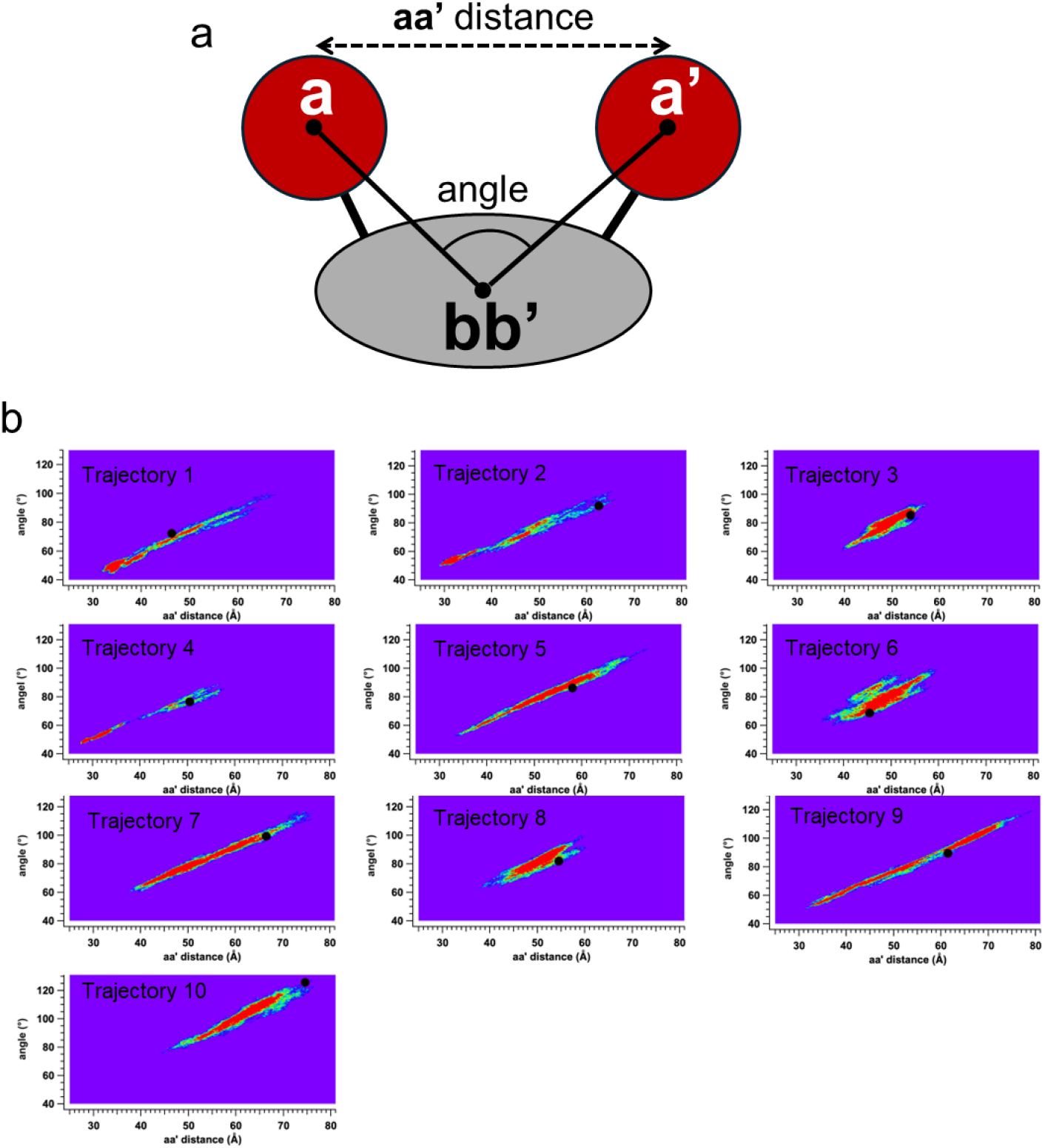
Conformational distributions derived from 10 independent MD trajectories. (a) Black circles denote domain centers of mass. Distance is defined between the **a** and **a’** domain centers, and angle as that formed by the **a**–**bb’**–**a’** centers. These parameters were used to compute conformational distributions for each trajectory. (b) 2D distributions of **a**–**a’** distance and angle for trajectories 1–10; black circle marks the first data point from MD production run.

Snapshots were extracted at 1-ps intervals from each trajectory, yielding a total of 100,001 structural conformations. For each structure, theoretical SAXS and SANS profiles of ER-60, as well as the SANS profile of segmentally deuterated ER-60, were calculated using Pepsi-SAXS or Pepsi-SANS. Averaging over all 100,001 structures yielded ensemble-averaged SAXS and SANS profiles. The same procedure was applied separately to each trajectory, resulting in a total of 30 averaged scattering profiles (10 trajectories × 3 experimental conditions). Using these averaged profiles, χ^2^ values were calculated against SAXS profile of *h*(WT), SANS profile of *h*(WT), and SANS profile of *h*(**a**)–*pd*(**bb’**)–*h*(**a’**) (Figure 6a-c). The trajectory that yielded the lowest χ^2^ value differed depending on whether SAXS profile of *h*(WT) alone, SANS profile of *h*(WT) alone, or SANS profile of *h*(**a**)–*pd*(**bb’**)–*h*(**a’**) alone was considered (refer to solid arrows in Figure 6a-c). This observation indicates that selecting the optimal trajectory based solely on a single scattering profile may not be appropriate. We therefore evaluated the total χ^2^, defined as the sum of the χ^2^ values obtained for the three experimental datasets (Figure 6d). Trajectory 7 yielded the lowest total χ^2^ value and was thus considered the most consistent with the combined experimental constraints within the scope of the present test simulations. A comparison between the ensemble-averaged scattering profiles derived from trajectory 7 and the experimental data is shown in Figure 7. Although the experimental curves are qualitatively reproduced, the χ^2^ values indicate that further refinement remains necessary. These results suggest that more extensive MD simulations, including longer sampling times and force-field optimization, are required to adequately capture the real conformational ensemble of highly dynamic MDPs such as ER-60. Nevertheless, the present proof-of-concept analysis demonstrates that incorporation of domain-selective SANS data as complementary constraints alongside SAXS allows effective discrimination among multiple MD trajectories. In other words, the integrated strategy combining SAXS, segmentally deuterated iCM-SANS, and MD simulations constitutes a viable protocol for high-precision dynamical analysis of MDPs. This protocol is expected to be broadly applicable to other MDPs with more complex architectures.

**Figure 6.**
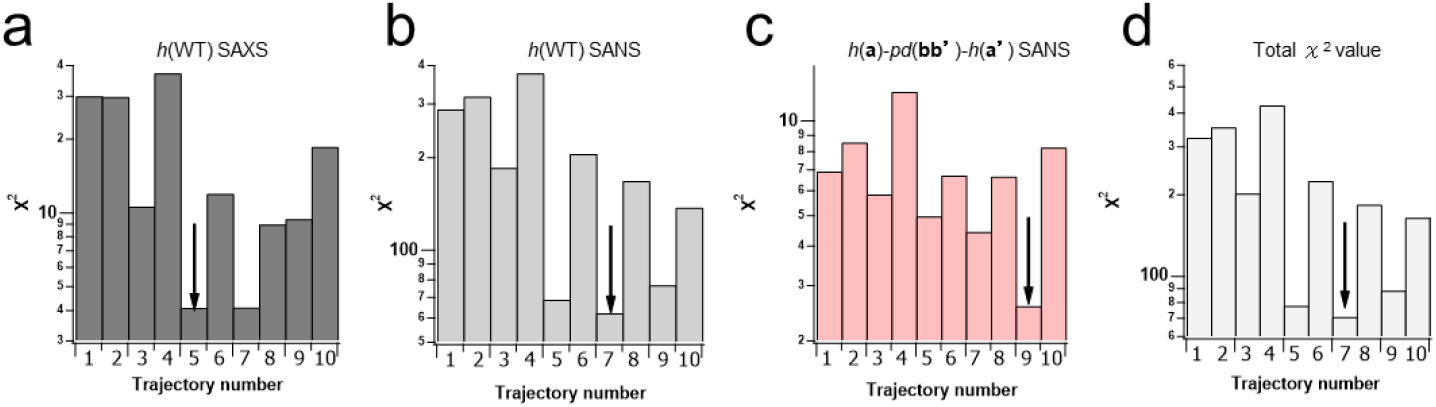
χ^2^ values of independent MD trajectories against SAXS, SANS of *h*(WT) and SANS of *h*(**a**)– *pd*(**bb’**)–*h*(**a’**). Arrows indicate the lowest χ^2^ values, corresponding to the best agreement with experimental profiles. (a–c) χ^2^ values for individual datasets: agreement between calculated scattering from each trajectory and experimental profiles of (a) SAXS of *h*(WT), (b) SANS of *h*(WT), and (c) SANS of *h*(**a**)–*pd*(**bb’**)–*h*(**a’**). (d) Total χ^2^, which is the addition of χ^2^ across all datasets.

**Figure 7.**
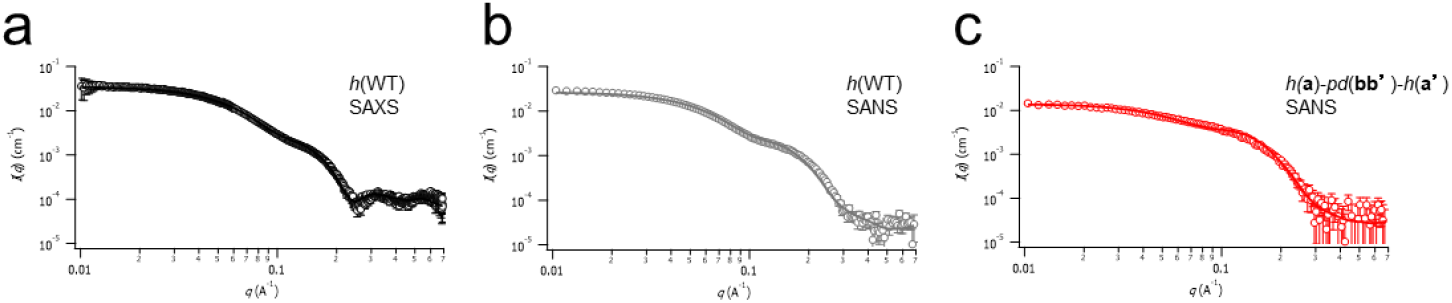
Comparison of experimental scattering profiles with ensemble-averaged theoretical profiles derived from Trajectory 7. (a) SAXS profile of *h*(WT), (b) SANS profile of *h*(WT), and (c) SANS profile of *h*(**a**)–*pd*(**bb’**)–*h*(**a’**). Experimental data (open symbols) are overlaid with ensemble-averaged theoretical profiles (solid lines) calculated from Trajectory 7.

## 4. Summary

In this study, we developed an integrated protocol for high-precision analysis of MDP dynamics by combining segmental deuteration attained by high-efficiency multi-step protein ligation, iCM-SANS, SAXS, and MD simulations. Using ER-60 as a model MDP, we successfully prepared segmentally deuterated constructs that enable selective visualization of spatially separated, non-contiguous domains without perturbing the native solution structure. Incorporation of domain-selective SANS data alongside SAXS enabled effective discrimination among independent MD trajectories, demonstrating that complementary scattering constraints are essential for resolving degeneracy in ensemble analysis of highly dynamic MDPs. Although extended simulations and further refinement will be required for fully quantitative ensemble determination, the present protocol supports a viable framework for integrating complementary scattering experiments with MD simulations. Importantly, this strategy is broadly applicable to other complex MDPs in which long-range inter-domain dynamics govern function. Looking forward, integration of this approach with high-resolution techniques such as NMR, which provides residue-level dynamical information, may further enhance ensemble discrimination and enable a more comprehensive description of domain–domain correlations of MDP in solution. Such multimodal integration represents a promising direction toward quantitative and mechanistic understanding of complex protein dynamics.

## Acknowledgements

SANS measurements were conducted with the approval of the Institut Laue-Langevin (ILL), and the Oak Ridge National Laboratory (ORNL) under proposal Nos. EASY-1099 and IPTS-31489.1. SAXS, AUC and MS measurements were conducted at KURNS under proposal Nos. R6113 and R7147.

## Data availability

The data that support the findings of this study are available from the corresponding author upon reasonable request.

