## Supporting information for "Integrating Segmental Deuteration iCM-SANS with SAXS and MD for Dynamical Analysis of Multi-domain Proteins"

### S1. Ligation efficiencies for the two-step ligation of segmentally deuterated ER-60.

**Table S1.** In addition to the efficiencies obtained in the present study, values reported in our previous work (Okuda *et al.*, 2023) are included for comparison.

| | Step | Ligation product | Ligation efficiency (%) |
|---|---|---|---|
| Present work | 1st | *pd*(**bb'**)-*h*(**a'**) | 62 |
| | 2nd | *h*(**a**)-*pd*(**bb'**)-*h*(**a'**) | 19 |
| Previous report [Okuda *et al.* 2023] | 1st | *h*(**bb'**)-*h*(**a'**) | 67 |
| | 2nd | *h*(**a**)-*h*(**bb'**)-*h*(**a'**) | 66 |

#### S2. Amino-acid sequence map of ER-60 used in segmental deuteration.

```
1   MRLRRLALFP GVALLLAAGR LVAASDVLEL TDDNFESRIS DTGSAGLMLV  50
51  EFFAPWCGHC KRLAPEYEAA ATRLKGIVPL AKVDCTANTN TCNKYGVSGY 100
                                  NGL
101 PTLKIFRDGE EAGAYDGPRT ADGIVSHLKK QAGPASVPLR TEEEFKKFIS 150
151 DKDASIVGFF DDSFSEAHSE FLKAASNLRD NYRFAHTNVE SLVNEYDDNG 200
201 EGIILFRPSH LTNKFEDKTV AYTEQKMTSG KIKKFIQENI FGICPHMTED 250
251 NKDLIQGKDL LIAYYDVDYE KNAKGSNYWR NRVMMVAKKF LDAGHKLNFA 300
301 VASRKTFSHE LSDFGLESTA GEIPVVAIRT AKGEKFVMQE EFSRDGKALE 350
                 NGL
351 RFLQDYFDGN LKRYLKSEPI PESNDGPVKV VVAENFDEIV NNENKDVLIE 400
401 FYAPWCGHCK NLEPKYKELG EKLSKDPNIV IAKMDATAND VPSPYEVRGF 450
451 PTIYFSPANK KLNPKKYEGG RELSDFISYL QREATNPPVI QEEKPKKKKK 500
501 AQEDL 505
```

**Figure S1.** Primary sequence of ER-60 showing hydrogenated and deuterated regions and ligation sites. Residues in red correspond to hydrogenated regions comprising the **a** and **a’** domains, whereas residues in gray correspond to deuterated regions comprising the **bb’** domains. NGL sequences indicate the *Oa*AEP recognition sites introduced for the two-step ligation. Cysteine residues are highlighted in blue. Catalytic CGHC motifs are shown in bold.

**S3. Mass spectra of hydrogenated and partially deuterated His-Smt3-bb' domains.**

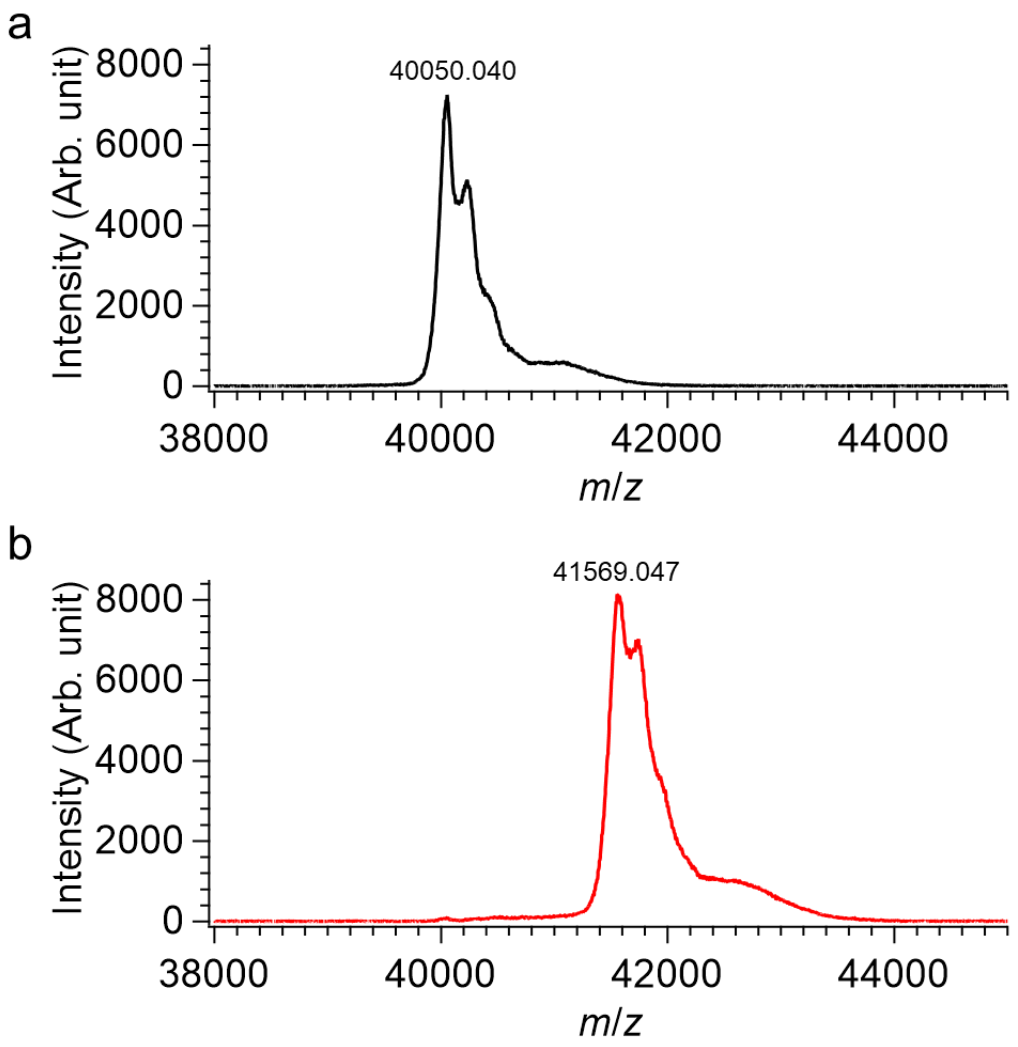


**Figure S2.** MALDI-TOF mass spectra of His-Smt3-**bb'** domain expressed under hydrogenated (a) and partially deuterated (b) conditions.

**S4. AUC profiles of ER-60 constructs.**

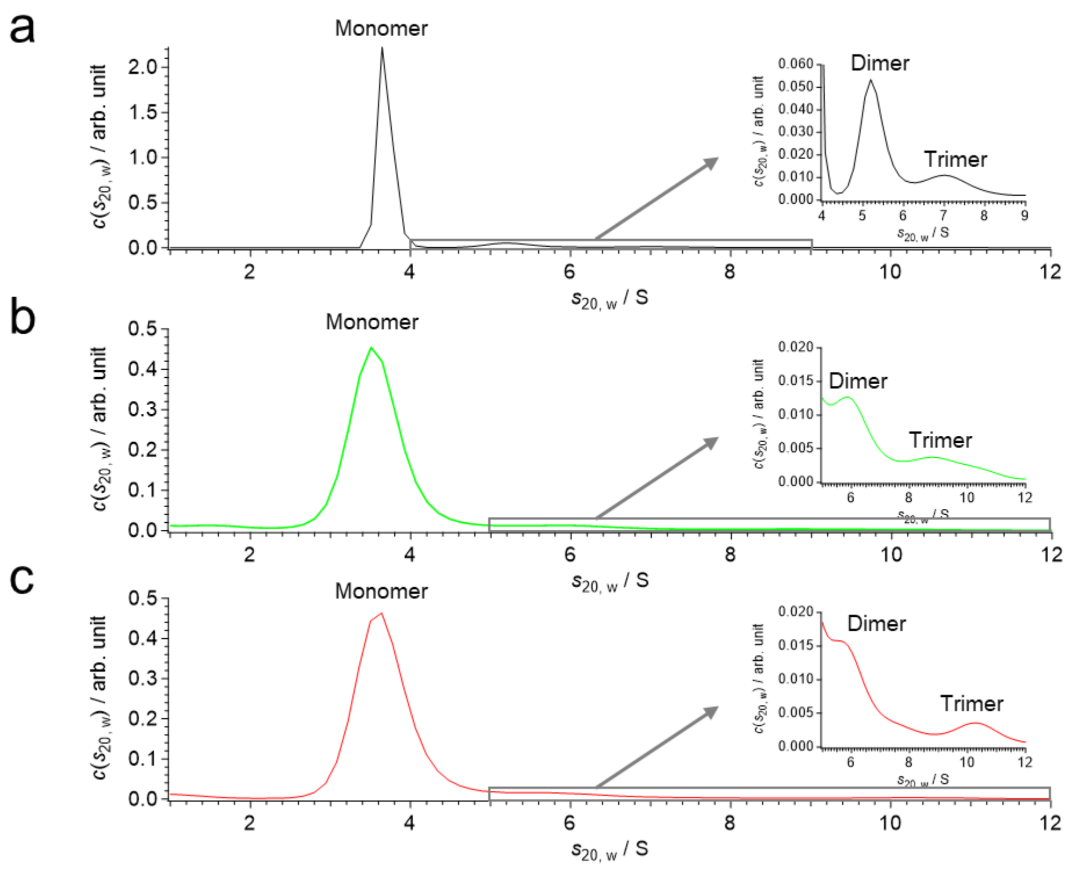


**Figure S3.** Sedimentation coefficient distributions of (a) *h*(WT), (b) *h*(**a**–**bb'**–**a'**), and (c) *h*(**a**)–*pd*(**bb'**)–*h*(**a'**). In all samples, the major peak corresponds to the monomer, with minor dimer and trimer species. Insets show magnified views of the high molecular components.

**S5. Sedimentation coefficients ($s_{20,w}$) and weight fractions of monomeric, dimeric, and trimeric species of *h*(WT), *h*(a-bb'-a'), and *h*(a)-*pd*(bb')-*h*(a').**

**Table S2.** Sedimentation coefficients ($s_{20,w}$) and weight fractions of monomeric, dimeric, and trimeric species of *h*(WT), *h*(**a-bb'-a'**), and *h*(**a**)-*pd*(**bb'**)-*h*(**a'**).

| | Monomer | | Dimer | | Trimer | |
|---|---|---|---|---|---|---|
| | $s_{20,w}$ (s) | Weight fraction (%) | $s_{20,w}$ (s) | Weight fraction (%) | $s_{20,w}$ (s) | Weight fraction (%) |
| *h*(WT) | 3.6 | 89.5 | 5.2 | 7.0 | 7.0 | 3.5 |
| *h*(**a-bb'-a'**) | 3.5 | 93.4 | 5.9 | 5.2 | 9.1 | 1.4 |
| *h*(**a**)-*pd*(**bb'**)-*h*(**a'**) | 3.6 | 91.7 | 4.9 | 6.9 | 7.0 | 1.4 |

**S6. SEC chromatogram and time course of integrated intensity of ER-60 constructs.**

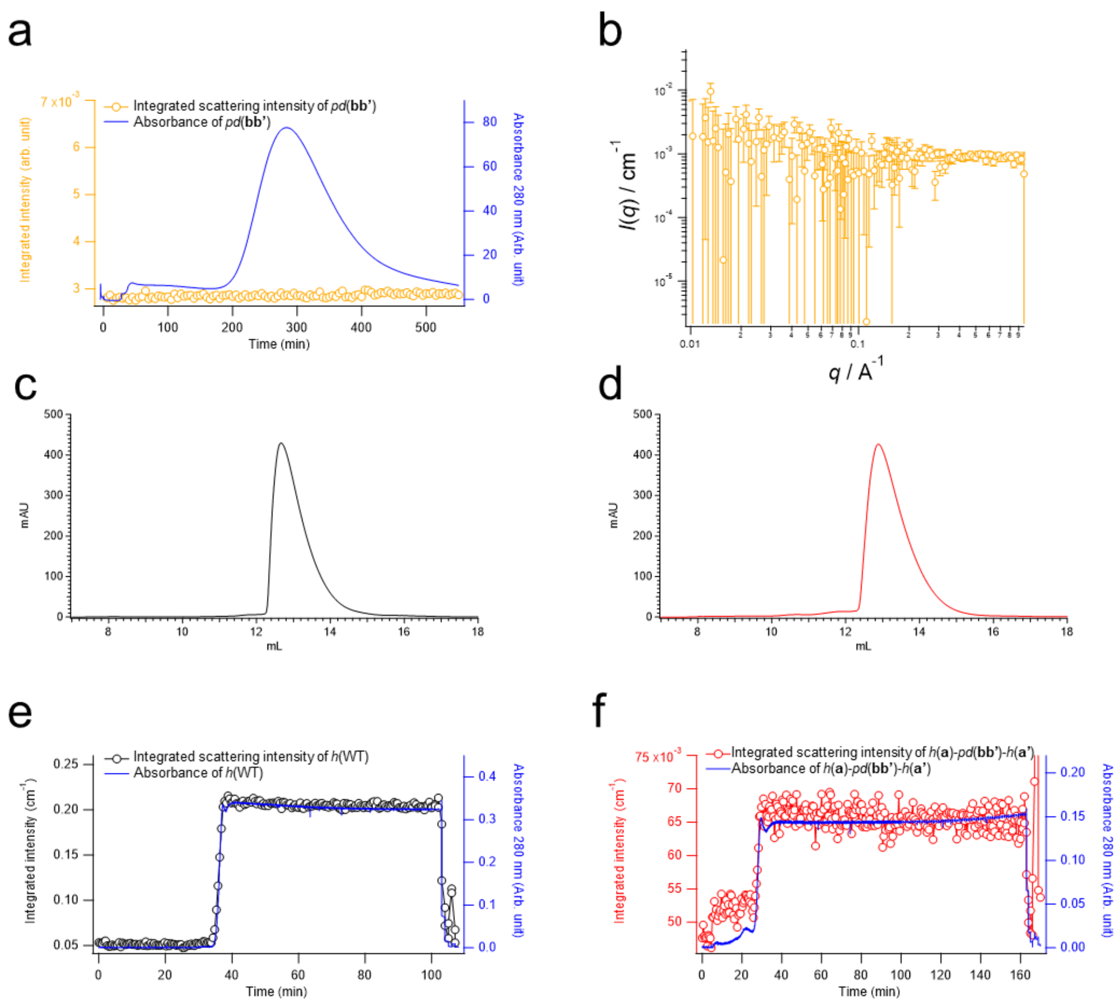


**Figure S4.** (a) SEC-SANS profile of deuterated **bb'** domain (*pd*(**bb'**)): The scattering intensity $I(_q)$ vs $q$ plot for *pd*(**bb'**).

(a) Time course of integrated intensity over the $q$ range 0.01–0.04 Å$^{-1}$ (open circles) and UV absorbance at 280 nm (blue line) of *pd*(**bb'**).

(b) SANS profile of *pd*(**bb'**).

(c) SEC elution chart of hydrogenated wild-type ER-60 (*h*(WT)).

(d) SEC elution chart of segmentally deuterated ER-60 (*h*(**a**)-*pd*(**bb'**)-*h*(**a'**))

(e) Time course of i integrated intensity over the $q$ range 0.01–0.03 Å$^{-1}$ (open circles) and UV absorbance at 280 nm (blue line) of *h*(WT).

(f) Time course of i integrated intensity over the $q$ range 0.01–0.03 Å$^{-1}$ (open circles) and UV absorbance at 280 nm (blue line) of *h*(**a**)-*pd*(**bb'**)-*h*(**a'**).
